# A deep learning model trained on only eight whole-slide images accurately segments tumors: wise data use versus big data

**DOI:** 10.1101/2022.02.07.478680

**Authors:** T. Perennec, R. Bourgade, Sébastien Henno, Christine Sagan, Claire Toquet, N. Rioux-Leclercq, Solène-Florence Kammerer-Jacquet, D. Loussouarn, M. Griebel

**Author notes:** these authors contributed equally to this work and should be regarded as joint first authors / last authors (T. Perennec, R. Bourgade, D. Loussouarn, M. Griebel). Corresponding author: T. Perennec.

## Abstract

Computer-assisted pathology is one of the biggest challenges in the medicine of the future. However, artificial intelligence is struggling to gain acceptance in the broader medical community due to data security issues, lack of trust in the machine, and poor data availability. Here, we develop a tumor delineation algorithm with only eight whole slide images of ovarian cancer to demonstrate the feasibility of an artificial intelligence application created from only a few data, finely annotated and with optimal processing. We test the model on seventeen other slides from the same hospital. The predictions are similar to the ground truth annotations made by an expert pathologist, with a mean DICE score of 0.90 [0.85 - 0.93]. The results on slides from another hospital are consistent, suggesting that the model is generalizable and that its performance does not suffer from different data acquisition. This study demonstrates the feasibility of a contouring algorithm based on a reduced dataset well optimized, going against the commonly accepted idea that a phenomenal amount of data is paramount. This study paves the way for other medical applications, especially for rare pathologies with limited available data.

## Introduction

In recent years, the amount of health data available has increased exponentially, both in volume and complexity, with the continuous arrival of new types of massive omics data (genomics, radiomics, proteomics, epigenomics, etc.). This phenomenon, called big data, has made the old dream of precision and personalized medicine a near prospect and a goal to achieve^1,2^. Several authors have pointed out the crucial role of data since the accuracy of machine learning and deep learning algorithms increases with the quantity and quality of data, but not significantly with the sophistication of the algorithms^3^. However, big data involves risks (patient identifications, theft of personal data, ethical concerns) and high costs (digitization, storage, energy, and computing time), so low and middle-income countries may miss this revolution, at the expense of the patient^4,5^. Several approaches allow for better data security, such as encryption or decentralized data with federating learning^6^, but reducing the amount of data used is the most obvious way to limit risk exposure.

Ovarian cancer represents 3.4 percent of cancers in women, with approximately 290 000 diagnoses worldwide each year. It is the eighth-most deadliest cancer in women^7^. Malignant epithelial tumors are the most common type of ovarian cancer, accounting for 90% of cases^8^. Like any cancer, histopathological analysis is the gold standard for diagnosing ovarian cancer. It allows the determination of at least five subtypes: high-grade serous carcinoma (70%), endometrioid carcinoma (10%), clear cell carcinoma (10%), mucinous carcinoma (3%), and low-grade serous carcinoma (<5%)^9^. It is crucial for the prognosis and the choice of treatment^10,11^ since these subtypes have different pathogenesis, molecular alterations, response to chemotherapy, and prognosis^10,12–14^. The ovarian cancer treatment landscape dramatically changed in 2014, with the arrival of PARP inhibitors in high-grade serous carcinomas^15,16^.

Deep learning is one of the most powerful AI methods in computer vision and is already widely studied in medical imaging and pathology^17^. It is a machine learning method based on a specific neural network for image processing called convolutional neural networks (CNN). Models learn the relationship between input, like an image, and its output, such as cancer location, by minimizing a loss function. The loss function represents the error of the prediction made by the model. It is minimized at each iteration by adjusting the internal parameters during back-propagation. This method can discover complex structures in large datasets, where conventional machine learning algorithms typically fail due to their limited ability to process raw data ^18^. The fields of application of deep learning in medicine are vast and allow the management of mass data in radiology, genomics, molecular biology, and information extraction in medical records^19–21^. However, computer-aided pathology must overcome additional challenges compared to radiology, such as the size and density of information contained in a single element and the usual lack of data availability^22,23^. Data from pathology, called whole-slide images (WSI), are obtained by digitizing histopathological slides thanks to the recent development of high-resolution scanners^24,25^.

Image segmentation is the process of determining which class each pixel belongs to (cancer or healthy tissue, for example). It has become a widely used tool in digital pathology research for the segmentation of small structures (cells, nuclei, or even glands) and, more rarely, entire regions^26^. Segmentation of nuclei or small repetitive structures like glands or glomerulus does not necessarily require large datasets because of their plurality in the whole slide image. For tumor segmentation, feature sizes vary (from emboli of a few micrometers to massive regions of several centimeters), making it challenging to identify small structures, especially if only massive structures are delineated in the training cohort and conversely. Tumor segmentation could save the pathologist time while increasing security by not missing any tumor spots. The predictions of segmentation models also have the advantage of being interpretable and therefore verifiable for a physician, unlike classification tools which only give the diagnosis in a WSI without further information.

Here, we develop and validate a deep learning model for the segmentation of ovarian cancer on whole-slide images based on a small annotated training dataset. In order to obtain a well-performing algorithm with a small amount of data, we use an innovative approach based on efficient sampling (figure 1).

**Figure 1.**
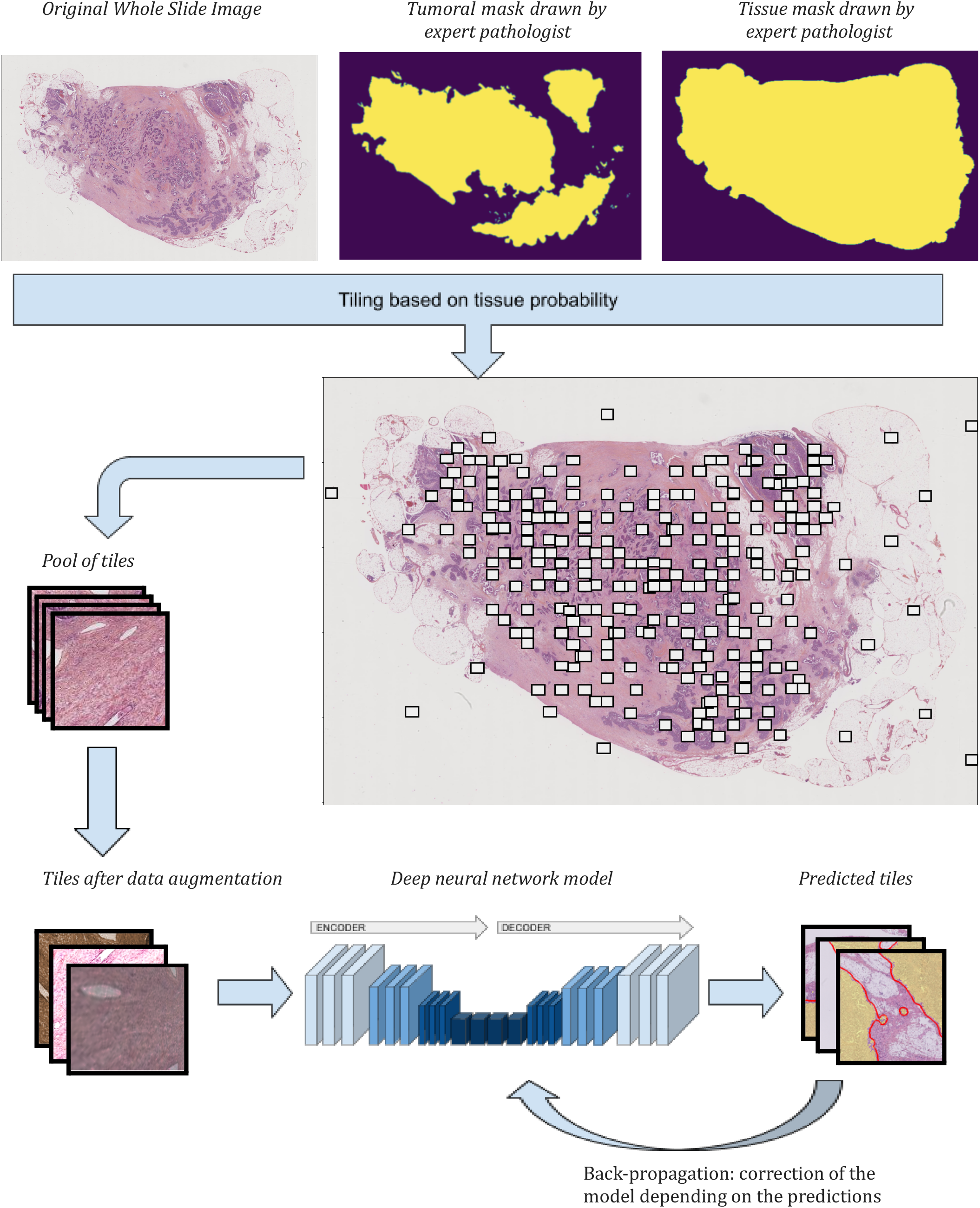
Complete workflow used for training the model: WSI and masks are tiled according to tissue and tumor probability before feeding the model.

## Results

### Internal performances

The average Dice score of the internal cohort is 0.898, and the mean pixel accuracy is 0.975. The Dice scores range from 0.529 to 0.984, 95% confidence interval of the mean obtained by bootstrap ranges from 0.85 to 0.93. Dice score, intersection over union (IoU), and pixel accuracy for each WSI of the validation cohort are presented in table 1. All of the predictions and the original WSI are available in supplementary data.

**Table 1.** Performances of the model over the internal validation cohort.

|  | Dice score | IoU | pixel accuracy |
| --- | --- | --- | --- |
| WSI 1 | 0.797 | 0.662 | 0.918 |
| WSI 2 | 0.931 | 0.872 | 0.987 |
| WSI 3 | 0.888 | 0.798 | 0.946 |
| WSI 4 | 0.925 | 0.861 | 0.990 |
| WSI 5 | 0.877 | 0.782 | 0.924 |
| WSI 6 | 0.910 | 0.835 | 0.964 |
| WSI 7 | 0.949 | 0.902 | 0.993 |
| WSI 8 | 0.940 | 0.886 | 0.992 |
| WSI 9 | 0.918 | 0.848 | 0.974 |
| WSI 10 | 0.859 | 0.753 | 0.993 |
| WSI 11 | 0.909 | 0.833 | 0.980 |
| WSI 12 | 0.960 | 0.922 | 0.981 |
| WSI 13 | 0.954 | 0.912 | 0.972 |
| WSI 14 | 0.984 | 0.969 | 0.985 |
| WSI 15 | 0.966 | 0.935 | 0.987 |
| WSI 16 | 0.963 | 0.928 | 0.989 |
| WSI 17 | 0.529 | 0.360 | 0.998 |
| **Mean** | **0.898** | **0.827** | **0.975** |

WSI 1 and 17 particularly stand out from the rest, with Dice scores below 0.8. Their image and their masks (predicted and annotated) are presented in Figure 2. In the first WSI, due to the papillary and micro-papillary architecture, the pathologists who segmented the images delineated several large tumor areas, including stroma and papillary lumen. In contrast, the algorithm was much more fine-grained in the segmentation, retaining only tumor pixels, as shown in figure 2A. On the other hand, small carcinomatous spots are missed by the algorithm on WSI 1, in the lower-left corner of the first slide, for instance (Figure 2 A). However, because these false-positive areas are small, they have little impact on the DICE score and even less on the IoU and pixel accuracy. All tumoral areas were delineated by the pathologist but some empty territories or consisting of stromal tissue have also been labeled. The inclusion of stromal tissue in the labeling is particularly unavoidable in tumor areas remodeled by chemotherapy, where the tumor appears dislocated as isolated cells caught in a scar stroma, preventing fine and individual labeling of these cells. In contrast, thanks to a pixel-by-pixel prediction, these empty territories are excluded by both the algorithm and post-processing methods in order to strictly retain tumor tissue.

**Figure 2.**
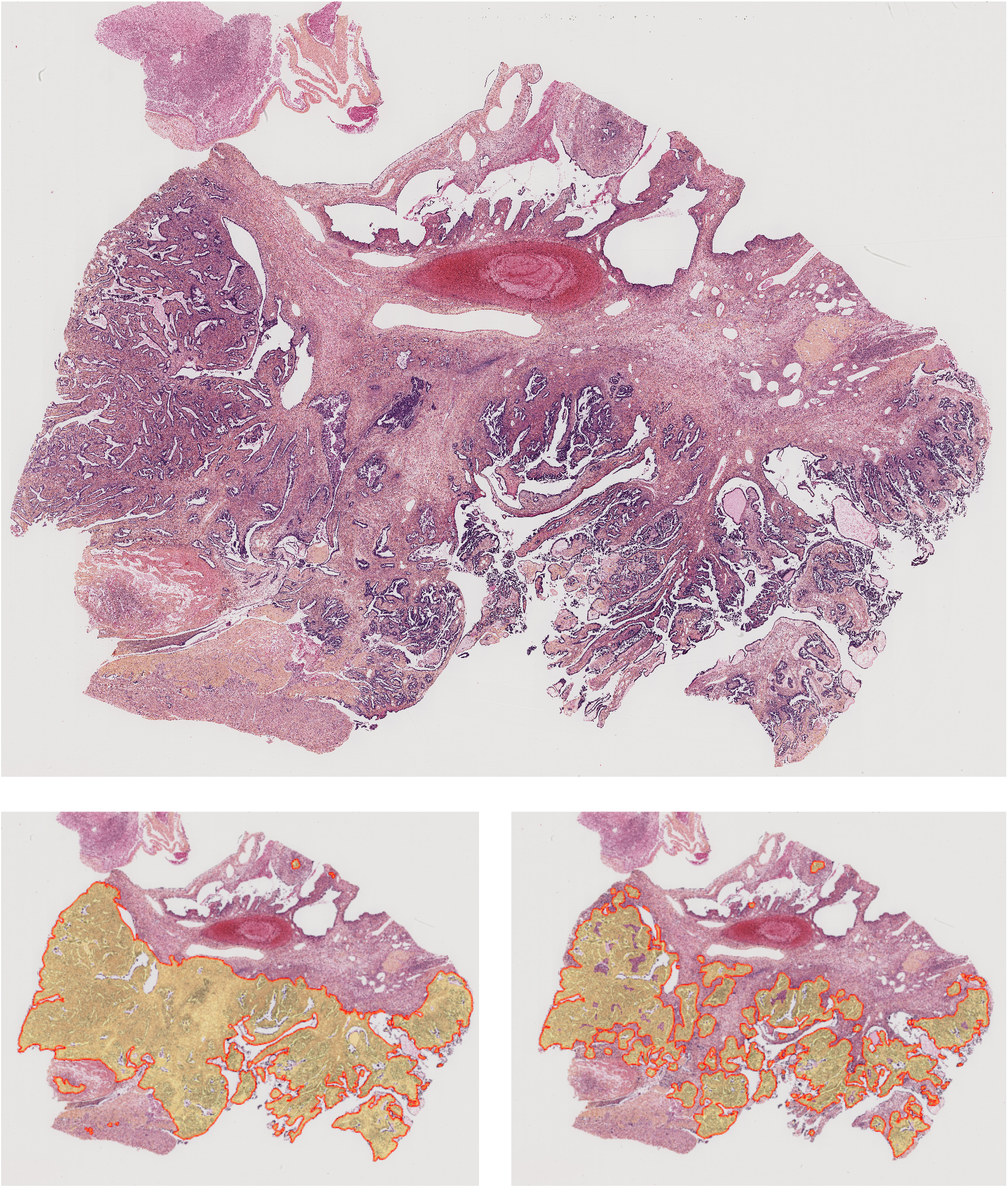
Image of the WSI, with the ground truth mask and the predicted mask for WSI 1 (A) and WSI 17 (B) *HES-stained slide: high-grade serous carcinoma with papillary and micropapillary architecture.* *A. WSI 1: image of the WSI, image of the WSI segmented by the pathologist (bottom left), and WSI segmented by the algorithm (bottom right)* *B. WSI 17: WSI segmented by the pathologist (bottom left) and WSI segmented by the algorithm (bottom right). The black box in the WSI image represent the region presented below in the segmented images*

**Figure 2B:**
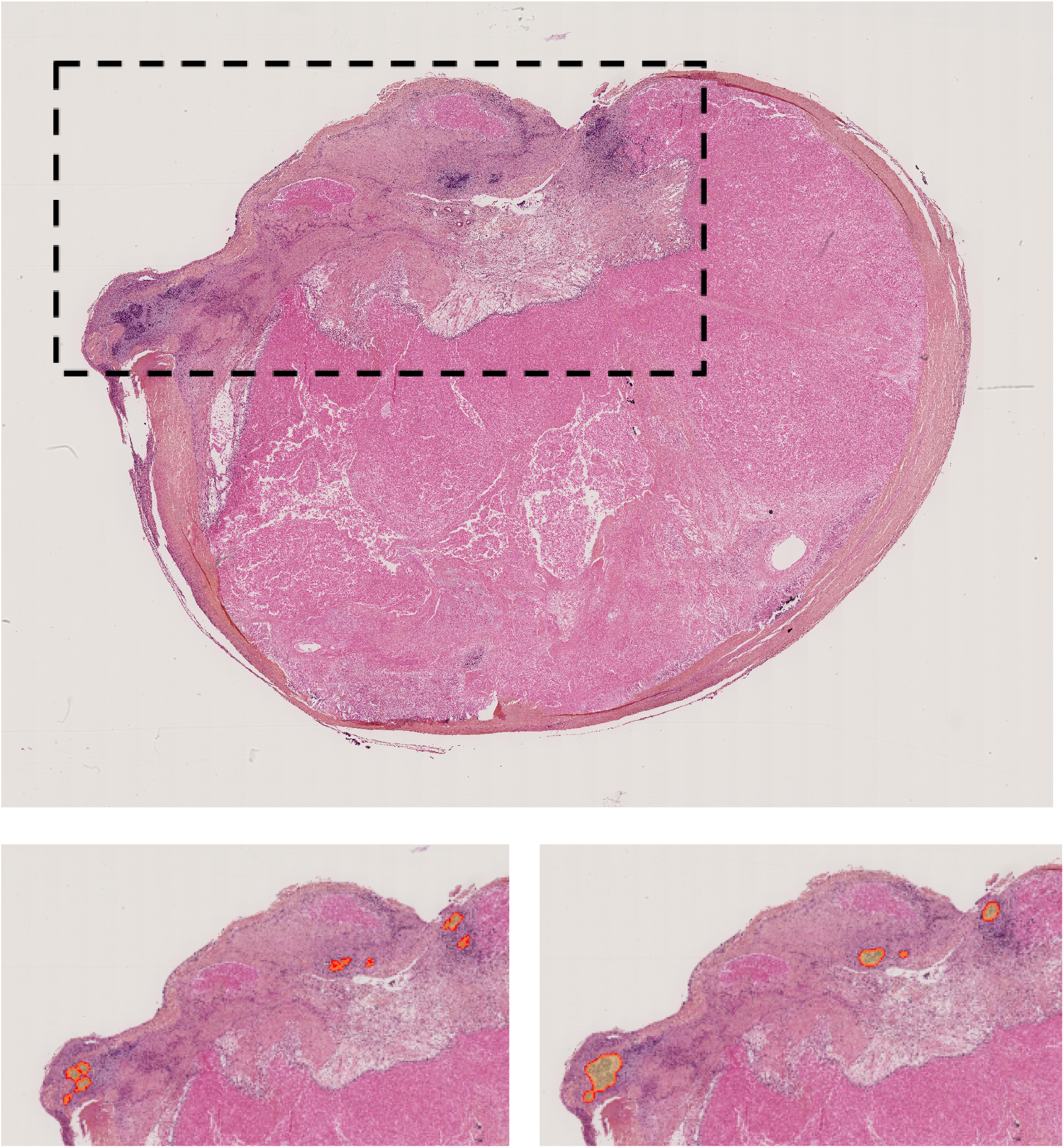
HES-staining lymph node with large area necrosis and a very few patches of carcinomatous cells.

WSI 17 is shown in Figure 2B. The tumor area is tiny compared to the rest of the WSI. Thus, although the prediction is very close to the ground truth, a few pixels difference is enough to significantly penalize the Dice and IoU scores, while the pixel accuracy is close to 1.

Artifacts related to the cutting formalin-fixed, paraffin-embedded tissue by microtome can lead to incorrect or inconclusive interpretations. Pathologists are trained to find artifacts and can differentiate artifacts from tissue components. The algorithm does not appear to be influenced by artifacts due to tissue folding, as shown in Figure 3. Such artifacts were not present in the training cohort.

**Figure 3.**
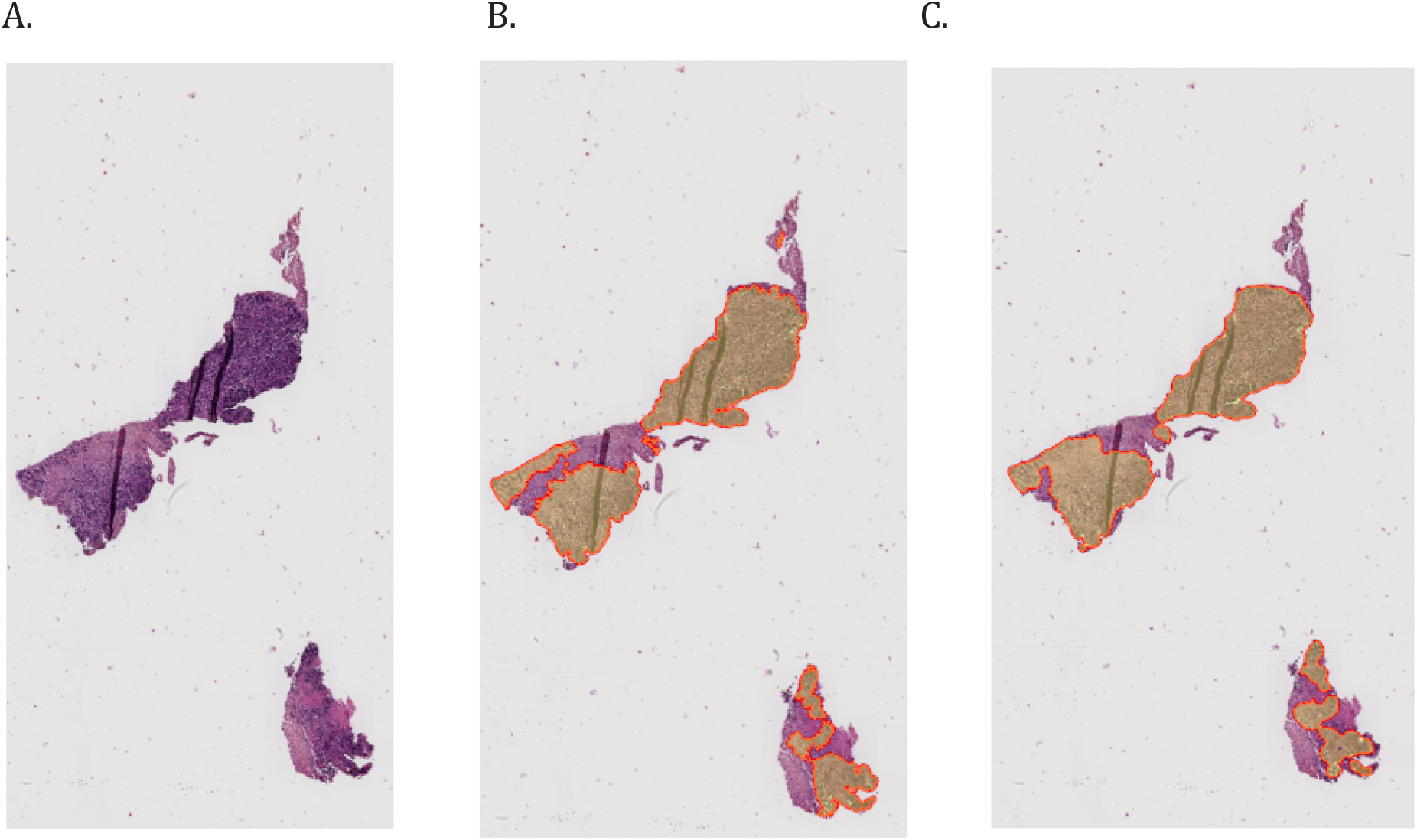
Images of predictions on a tissue sample with an artifact of WSI 4. *Image of the original WSI (A), WSI segmented by the pathologist (B), and WSI segmented by the algorithm (C)*

### External predictions

We made the predictions on an external cohort, from another hospital, with WSI numerized on another brand of scanner. A sample of the predictions is presented in figure 4. The entire set of predictions is available in the supplementary data. The predictions made by the algorithm were reviewed by a gynecopathologist expert who assessed that the majority of the carcinomatous areas were accurately delineated. Nevertheless, it missed some small tumoral areas. As with the internal dataset, predictions are more refined than humans could have made. For this reason, we did not calculate a score for the external cohort, preferring a visual comparison.

**Figure 4.**
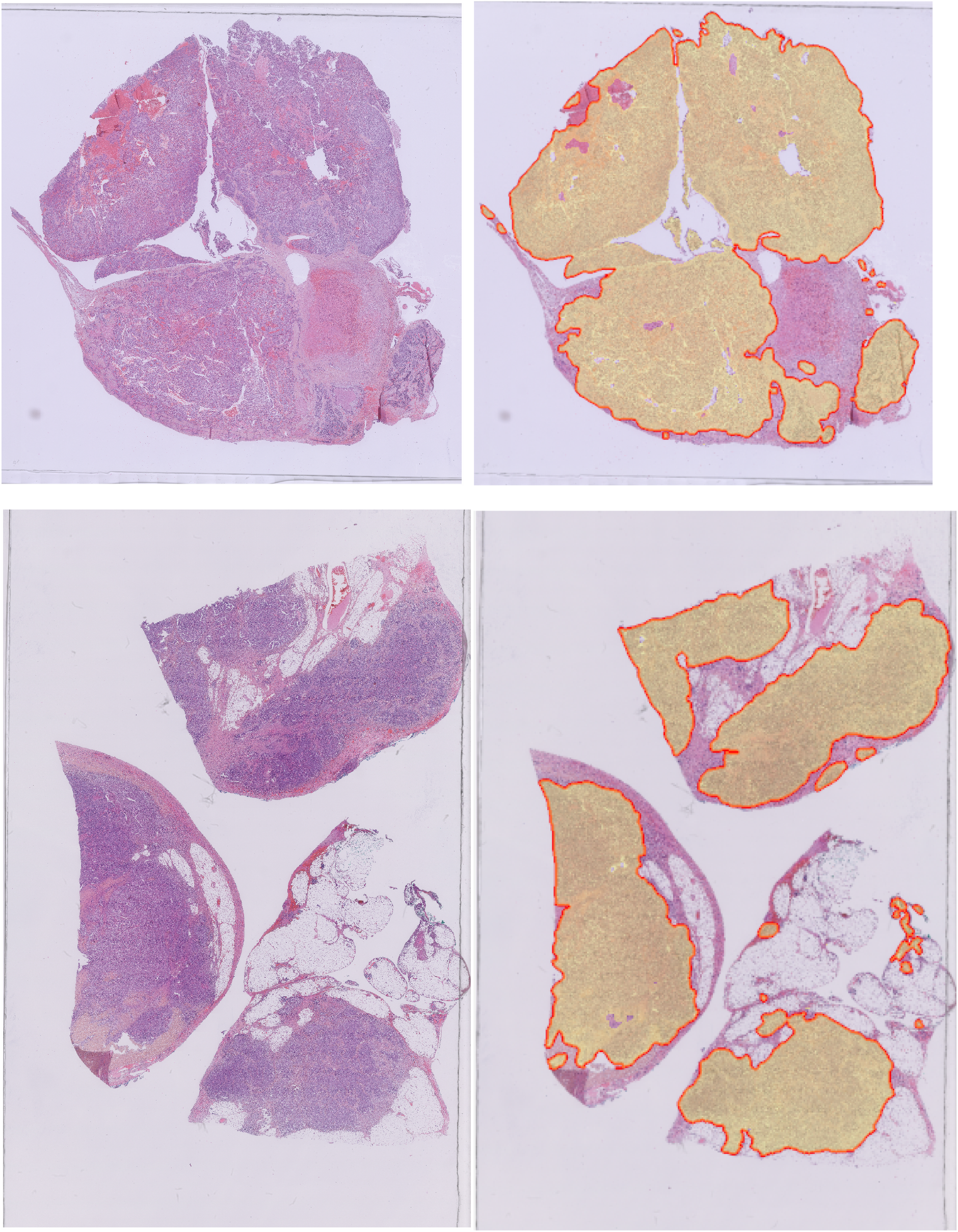
Predictions made on two WSI from the external dataset

## Discussion

Segmentation is a prerequisite for numerous pathology research projects. In deep learning projects, it can extract regions of interest to optimize learning by focusing on the relevant areas, like the tumor.

Deep learning usually requires a large amount of data, which is sometimes incompatible with rare diseases and current practice in many pathology departments that are under-equipped with high-resolution slide scanners and computing resources. Moreover, some countries will not be able to equip themselves for years to come, further widening the gap in research between wealthy and emerging countries and leaving out specific diseases of these populations. In addition, the correct labeling of WSI (to the nearest pixel) is very time-consuming and challenging for human pathologists, making the constitution of a quality annotated database difficult. Moreover, if we consider deep learning as assistance to delineation, the number of WSI labeled by the pathologist must be significantly lower than the total dataset. Here, we showed that obtaining a correct segmentation algorithm with only eight WSI is possible. The contours are not perfect but are similar to those made by the pathologist. It could be re-imported into visualization software to be corrected if needed. Once updated, it could even be reused for a new round of training. The learning process would be done in several steps, with a human-machine collaboration.

Although expert pathologists can give a qualitative assessment, an accurate and reproducible assessment by a widely established metric would ideally be preferred. The evaluation of the model is made difficult here by the absence of perfect ground truth. Such a gold standard is impossible to make for a human pathologist. Indeed, despite the careful contouring performed by the gynecopathologist expert, she has inevitably selected non-tumoral tissue. Therefore, a hypothetical perfect segmentation made by an algorithm (extraction of the whole tumor and no non-tumoral pixels) would not obtain a maximum score since the metric measures the difference between the contours drawn by humans and those drawn by the algorithm. Furthermore, a weak correlation between expert assessment and traditional metrics has been noted^27^. We used the DICE coefficient, the most used metric in biomedical image analysis competition^28^, even though there is no consensus on the method to evaluate the models. The lack of stability in the results (significant changes in results after minor changes in the data) and the critical changes induced by a metric or aggregation method (median or mean) calls for great caution in interpreting numerical results from standard scores^28^. In the absence of an effective method to properly evaluate the models, it should be kept in mind that such algorithms can only be considered as tools to assist delineation, but not substitutes. It has already been shown that a clinician assisted by AI performs better than a clinician alone in classification tasks^29,30^, and sometimes more than both human and AI^31^. In our case, the algorithm is an excellent tool to remove the non-tumoral tissue, necessarily included by the pathologist. On the other hand, a review by a pathologist is necessary to be sure not to miss any tumor spot.

The algorithm obtained in this study does not allow a perfect delineation of the tumor areas because it lacks sensitivity on some small tumor spots but outperforms human delineation on specificity. The results obtained are, to our knowledge, the only ones showing the delineation of large structures with so little data. These results call for a more reasoned use of data when the current trend is using big data at all costs.

## Methods

### Patients

The training and internal validation dataset are composed of eight and seventeen patients respectively, randomly selected from all the patients diagnosed with a high-grade ovarian carcinoma between January 2016 and December 2019 in the University Hospital of Nantes in France. To demonstrate the ability of the algorithm to generalize its results, we used an external validation dataset of eight patients selected from the pathology laboratory of the University Hospital of Rennes in France. All images were anonymized before the study, and no clinical data were collected.

### Pre-processing

All images used for training were HES-stained, Formalin-Fixed Paraffin-Embedded diagnostic sections from high-grade ovarian cancer. If several slides were available for a single patient, we selected for training the slide with visually the most tumor. The slides used for the training and the internal validation dataset were digitized at 40x magnification (0.22μm/px) using the Hamamatsu Nanozoomer scanner in the Micropicell platform (IRS-UN unit, Nantes). The external validation dataset consists of eight WSIs digitized using a Philips scanner at 40x magnification (0.25μm/px). The three datasets were anonymized and downscaled to 20x magnification (0.5μm/px) to optimize memory. All WSIs from training and validation datasets were manually annotated on Qupath (version 0.2.3) by one experienced gynecopathologist, drawing for each WSI one tumor mask and one mask surrounding all the tissue to optimize sampling. The polygonal annotations were then exported using the GeoJSON format and converted into a binary matrix used for supervised training.

All slides were normalized by rescaling pixel values from 0-255 to 0-1. The mean and the standard deviation used for normalization were computed on each channel of the dataset. Instead of tiling the entire WSI as usually done, we opted for a memory-efficient sampling strategy combining two approaches. We first sampled tiles of 512 pixels per side centered on tumor areas using the tumor mask and combined them with random tiles sampled according to a probability density function on the tissue mask. A hundred tiles per image and per epoch were thus sampled with a 0.9 probability for tissue and 0.1 for the background region to ensure that our model can interpret it as well.

All tiles were finally randomly transformed using affine transformations and color parameter variations. This data augmentation reduces the risk of overfitting by artificially increasing the dataset.

### Training

The pipeline has been realized under Python (version 3.7) using Pytorch framework and Scikit-learn, Fastai, and Deepflash2 libraries^32^. All calculations have been made on an NVIDIA V100 Tensor Core GPU with 32 Gb of memory. We used a fully convolutional network pre-trained on ImageNet with a U-Net architecture and an EfficientNet-B6 as a backbone to perform semantic segmentation. We used Ranger as optimizer and a cross-entropy loss. Ranger combines a rectified version of Adam (RAdam) and a LookAhead optimizer for a faster convergence that is less dependent on the learning rate. We trained the model in mixed precision to allow a faster forward propagation with a 0.01 weight decay. Twelve epochs with a batch size of 16 have been performed using the one-cycle policy with a 10-3 max learning rate.

### Post-processing

Post-processing includes all modifications made to the predictions of the algorithm. It consists of two steps. The first step is smoothing and consists in replacing each 12 by 12 pixel square with a new square containing only the 90th percentile value of the first square, as shown in figure 5. The second step consists in removing the pixels predicted as tumors from the areas containing no tissue. For this, we perform the intersection between the predicted mask and the initial tissue mask, automatically determined by thresholding on the blue color channel, as shown in figure 6.

**Figure 5.**
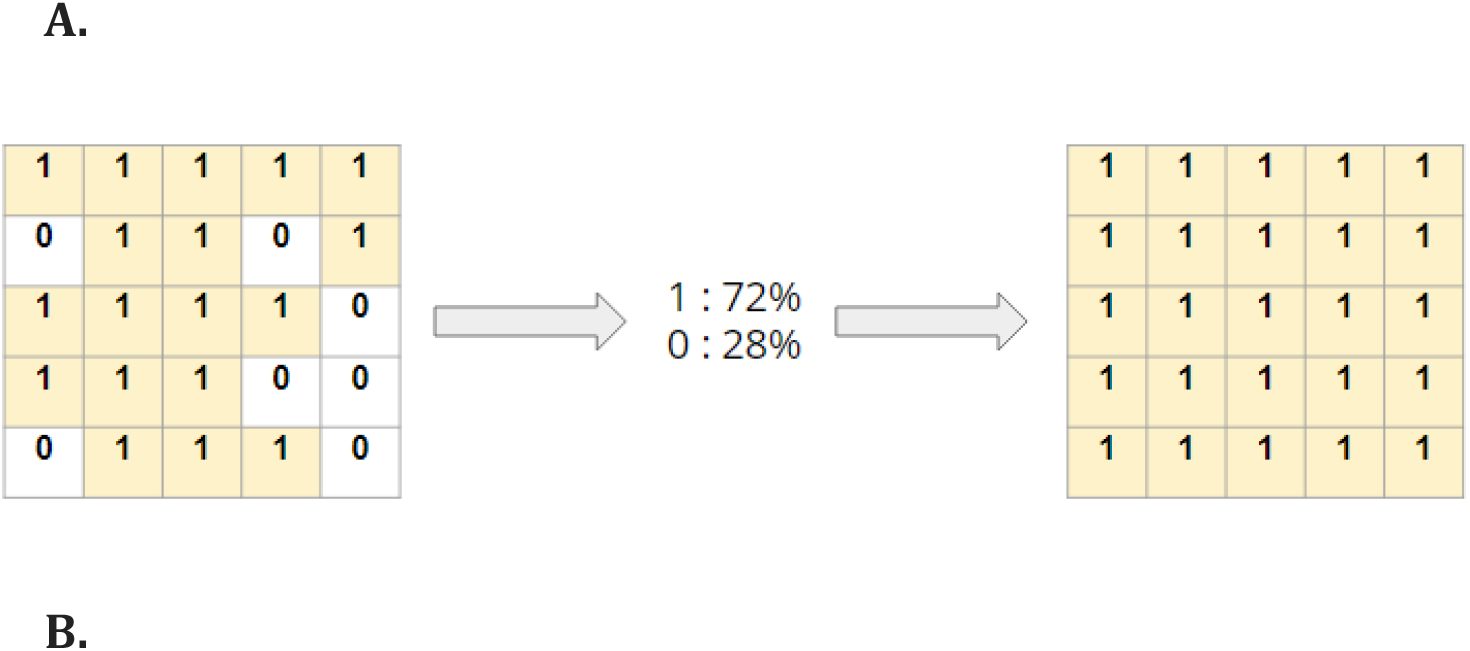
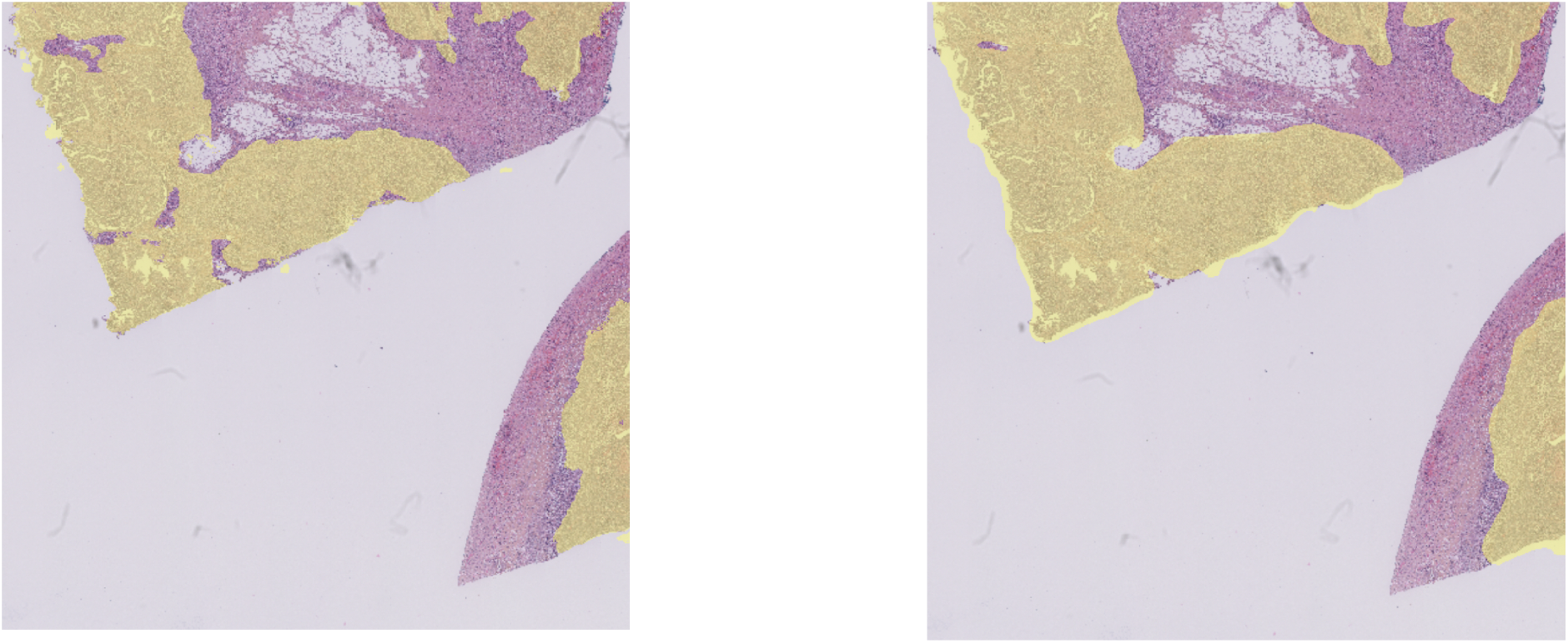
Schematic representation (A) and example (B) of the smoothing by the 90th percentile

**Figure 5.**
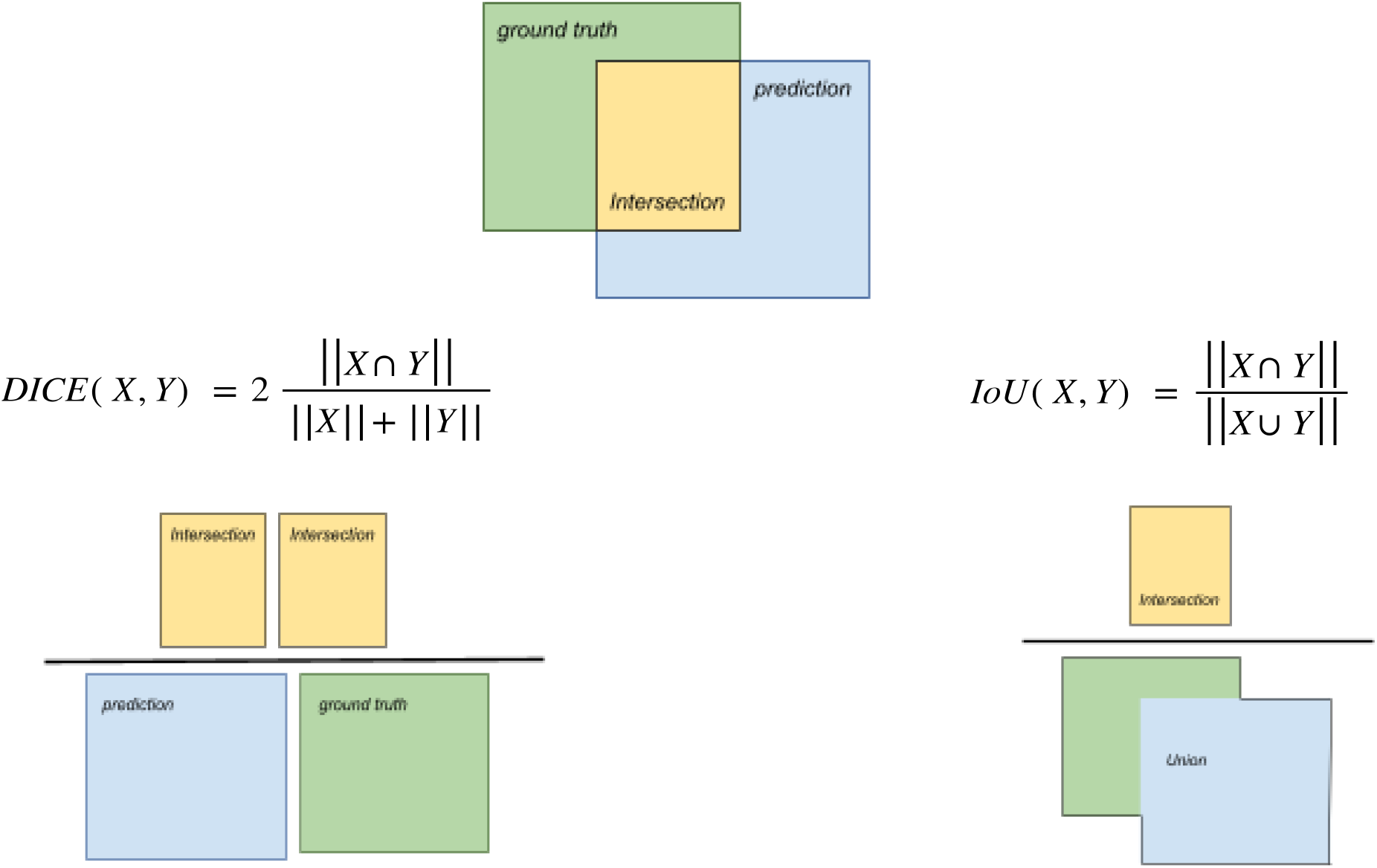
The formula of the Dice score and IoU and schematic representations of their definition. *X and Y are the two segmentation masks for the tumor class. ||X|| and ||Y|| are the norm (the area) of X and Y, and ||X* ⋂ *Y|| is the norm of their intersection.* ⋂ *and* ⋃ *are the intersection and union operators.*

**Figure 6.**
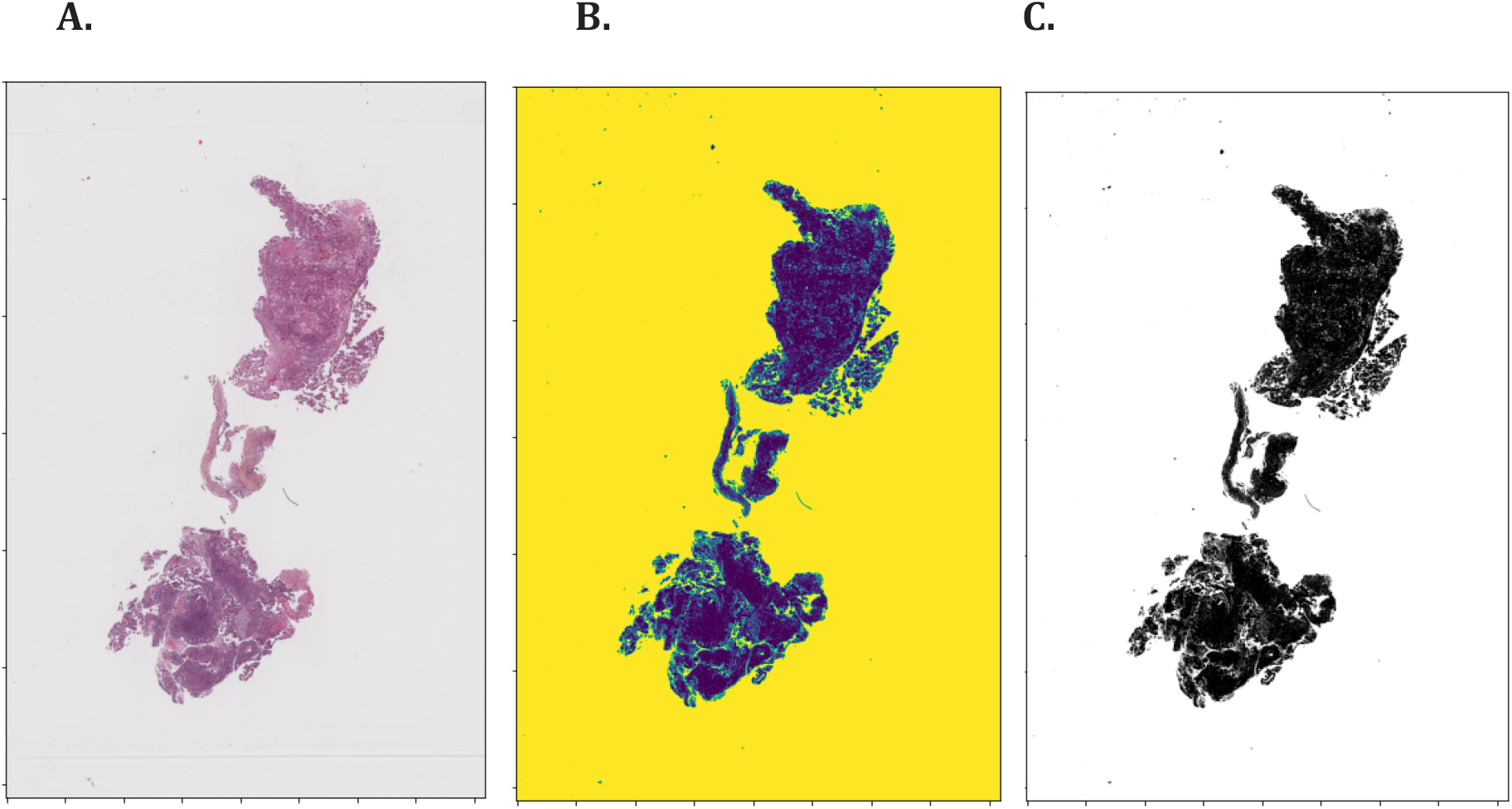
Example of tissue extraction by thresholding on blue color value. The original slide (A) is transformed to keep only the blue channel (B), and we then only extract the darkest pixels (C)

### Metrics and statistical analysis

The performance of the model was assessed by the Dice coefficient (or Sorensen-Dice index), the intersection over union (IoU), and the pixel accuracy. The Dice coefficient is defined as two times the intersection between the mask drawn by pathologists (called ground truth) and the prediction over the addition of the two surfaces. The formula and the schematic representation of each score are presented in figure 5.

## References

1. Ho, D. et al. Enabling Technologies for Personalized and Precision Medicine. Trends in Biotechnology 38, 497–518 (2020).

2. Hamburg, M. A. & Collins, F. S. The Path to Personalized Medicine. New England Journal of Medicine 363, 301–304 (2010).

3. Kim, H.-S., Kim, D.-J. & Yoon, K.-H. Medical Big Data Is Not Yet Available: Why We Need Realism Rather than Exaggeration. Endocrinol Metab (Seoul) 34, 349–354 (2019).

4. Strubell, E., Ganesh, A. & McCallum, A. Energy and Policy Considerations for Modern Deep Learning Research. AAAI 34, 13693–13696 (2020).

5. Alyass, A., Turcotte, M. & Meyre, D. From big data analysis to personalized medicine for all: challenges and opportunities. BMC Medical Genomics 8, 33 (2015).

6. Rajpurkar, P., Chen, E., Banerjee, O. & Topol, E. J. AI in health and medicine. Nat Med 28, 31–38 (2022).

7. Bray, F. et al. Global cancer statistics 2018: GLOBOCAN estimates of incidence and mortality worldwide for 36 cancers in 185 countries. CA: A Cancer Journal for Clinicians 68, 394–424 (2018).

8. IARC. WHO Classification of Tumours Editorial Board: WHO Classification of Tumours. vol. 4 (2020).

9. Gilks, C. B. & Prat, J. Ovarian carcinoma pathology and genetics: recent advances. Hum Pathol 40, 1213–1223 (2009).

10. Roett, M. A. & Evans, P. Ovarian Cancer: An Overview. AFP 80, 609–616 (2009).

11. Köbel, M. et al. Ovarian Carcinoma Subtypes Are Different Diseases: Implications for Biomarker Studies. PLOS Medicine 5, e232 (2008).

12. du Bois, A. et al. A randomized clinical trial of cisplatin/paclitaxel versus carboplatin/paclitaxel as first-line treatment of ovarian cancer. J Natl Cancer Inst 95, 1320–1329 (2003).

13. Ledermann, J. A. PARP inhibitors in ovarian cancer. Ann Oncol 27 Suppl 1, i40–i44 (2016).

14. Ledermann, J. A. Front-line therapy of advanced ovarian cancer: new approaches. Ann Oncol 28, viii46–viii50 (2017).

15. Ledermann, J. A. PARP inhibitors in ovarian cancer. Ann Oncol 27 Suppl 1, i40–i44 (2016).

16. Ledermann, J. A. Front-line therapy of advanced ovarian cancer: new approaches. Ann Oncol 28, viii46–viii50 (2017).

17. Song, H., Nguyen, A.-D., Gong, M. & Lee, S. A Review of Computer Vision Methods for Purpose on Computer-Aided Diagnosis. Journal of International Society for Simulation Surgery 3, 1–8 (2016).

18. LeCun, Y., Bengio, Y. & Hinton, G. Deep learning. Nature 521, 436–444 (2015).

19. Shen, D., Wu, G. & Suk, H.-I. Deep Learning in Medical Image Analysis. 30 (2017).

20. Koumakis, L. Deep learning models in genomics; are we there yet? Computational and Structural Biotechnology Journal 18, 1466–1473 (2020).

21. Ching, T. et al. Opportunities and obstacles for deep learning in biology and medicine. Journal of The Royal Society Interface 15, 20170387 (2018).

22. Gurcan, M. N. et al. Histopathological Image Analysis: A Review. IEEE Rev Biomed Eng 2, 147–171 (2009).

23. Xu, Y. et al. Large scale tissue histopathology image classification, segmentation, and visualization via deep convolutional activation features. BMC Bioinformatics 18, 281 (2017).

24. Pantanowitz, L. et al. Twenty Years of Digital Pathology: An Overview of the Road Travelled, What is on the Horizon, and the Emergence of Vendor-Neutral Archives. J Pathol Inform 9, 40 (2018).

25. Pantanowitz, L., Farahani, N. & Parwani, A. Whole slide imaging in pathology: advantages, limitations, and emerging perspectives. PLMI 23 (2015) doi:10.2147/PLMI.S59826.

26. Deng, S. et al. Deep learning in digital pathology image analysis: a survey. Front. Med. 14, 470–487 (2020).

27. Kofler, F. et al. Are we using appropriate segmentation metrics? Identifying correlates of human expert perception for CNN training beyond rolling the DICE coefficient. arXiv:2103.06205 [cs, eess] (2021).

28. Maier-Hein, L. et al. Why rankings of biomedical image analysis competitions should be interpreted with care. Nat Commun 9, 5217 (2018).

29. Park, A. et al. Deep Learning-Assisted Diagnosis of Cerebral Aneurysms Using the HeadXNet Model. JAMA Netw Open 2, e195600 (2019).

30. Kim, H.-E. et al. Changes in cancer detection and false-positive recall in mammography using artificial intelligence: a retrospective, multireader study. Lancet Digit Health 2, e138–e148 (2020).

31. Steiner, D. F. et al. Impact of Deep Learning Assistance on the Histopathologic Review of Lymph Nodes for Metastatic Breast Cancer. Am J Surg Pathol 42, 1636–1646 (2018).

32. Segebarth, D. et al. DeepFLaSH, a deep learning pipeline for segmentation of fluorescent labels in microscopy images. 473199 https://www.biorxiv.org/content/10.1101/473199v1 (2018) doi:10.1101/473199.

