## Supplementary figures for "A deep learning model trained on only eight whole-slide images accurately segments tumors: wise data use versus big data"

### Supplementary data

**Supplementary figure 1**. WSI of Nantes delineated by the pathologist and the algorithm

| WSI | WSI delineated by the pathologist | WSI delineated by the algorithm |
|---|---|---|
| 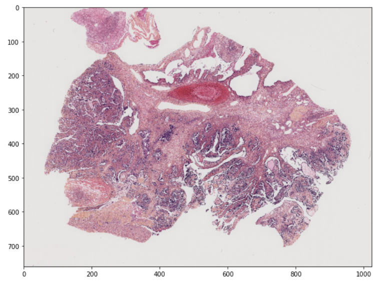 | 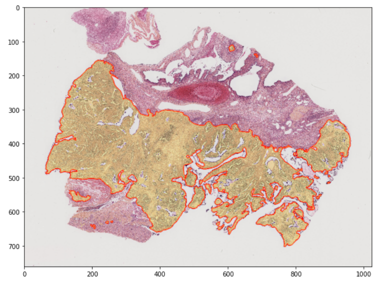 | 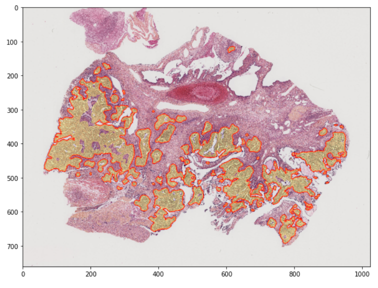 |
| 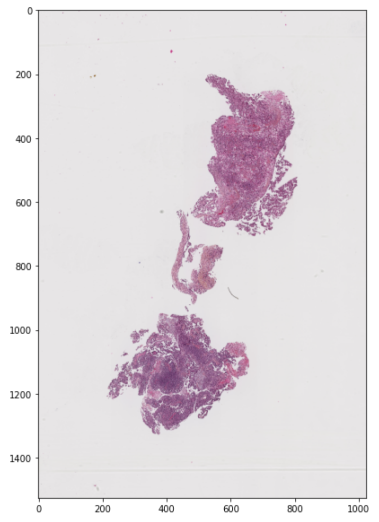 | 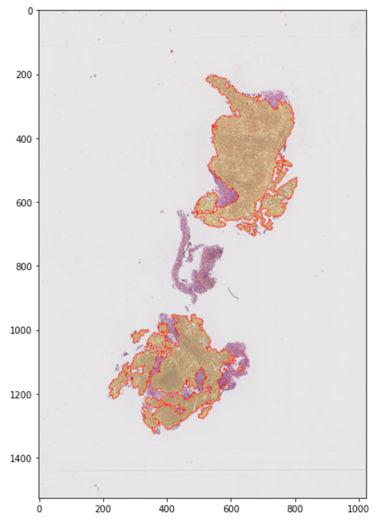 | 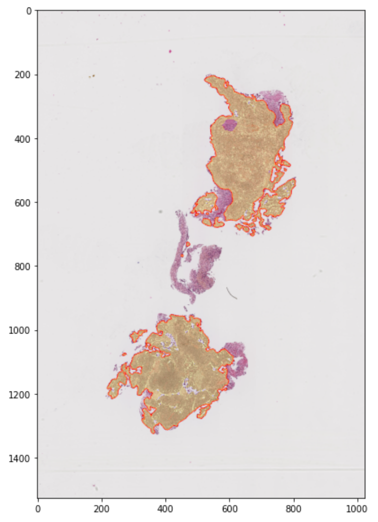 |
| 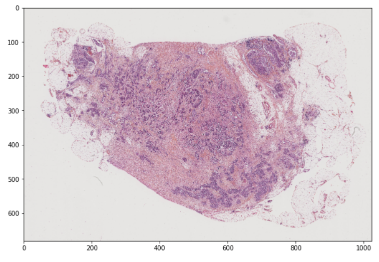 | 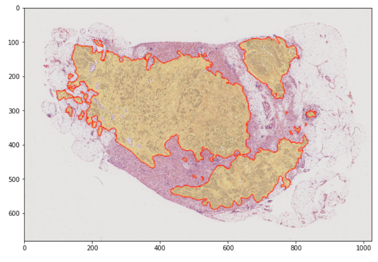 | 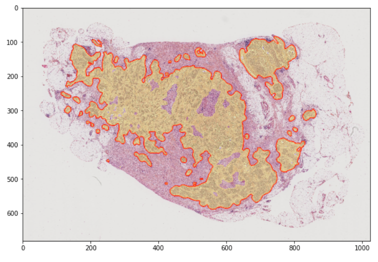 |

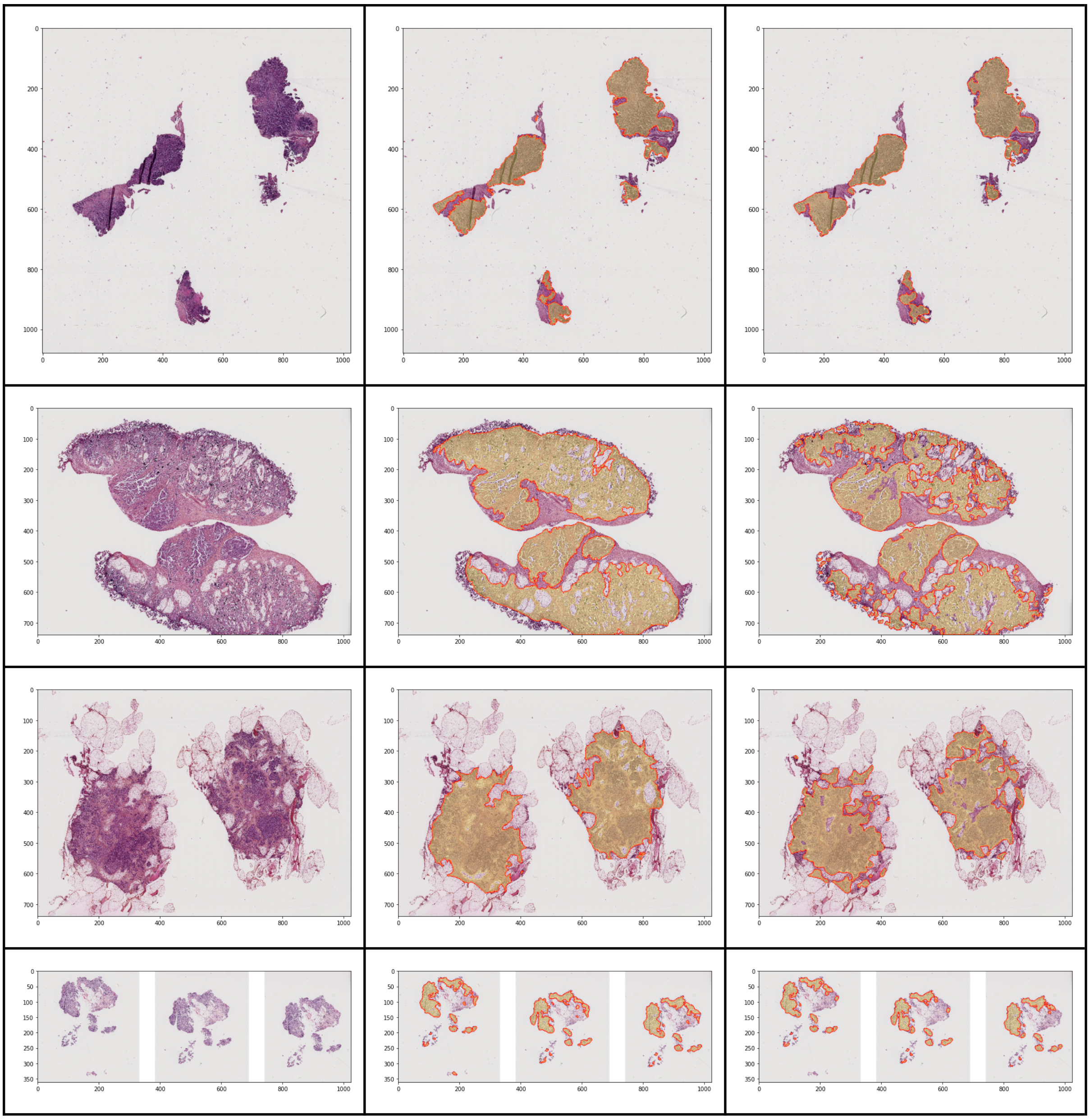

0
200
400
600
800
1000
100
300
500
700
50
150
250
350

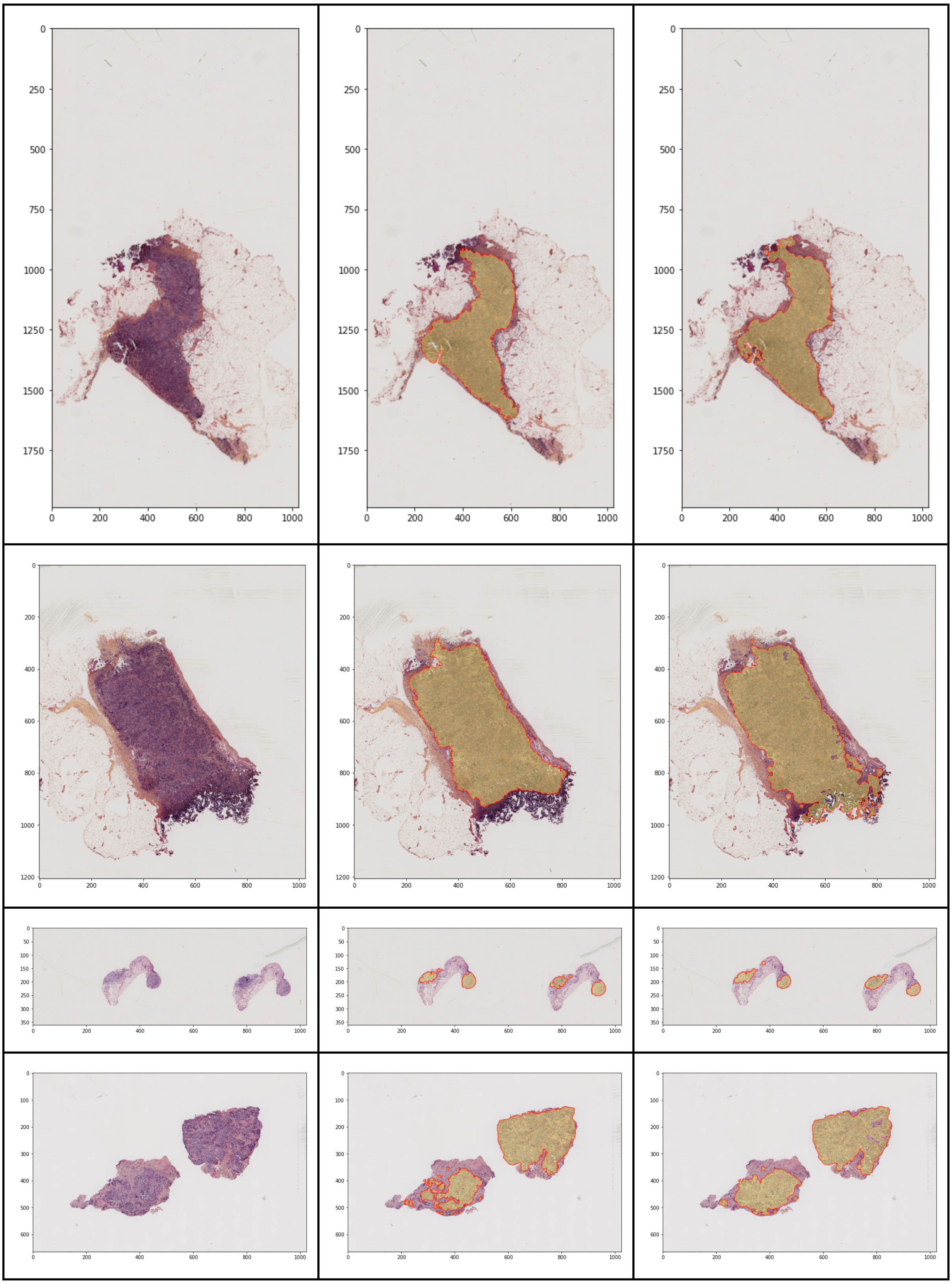

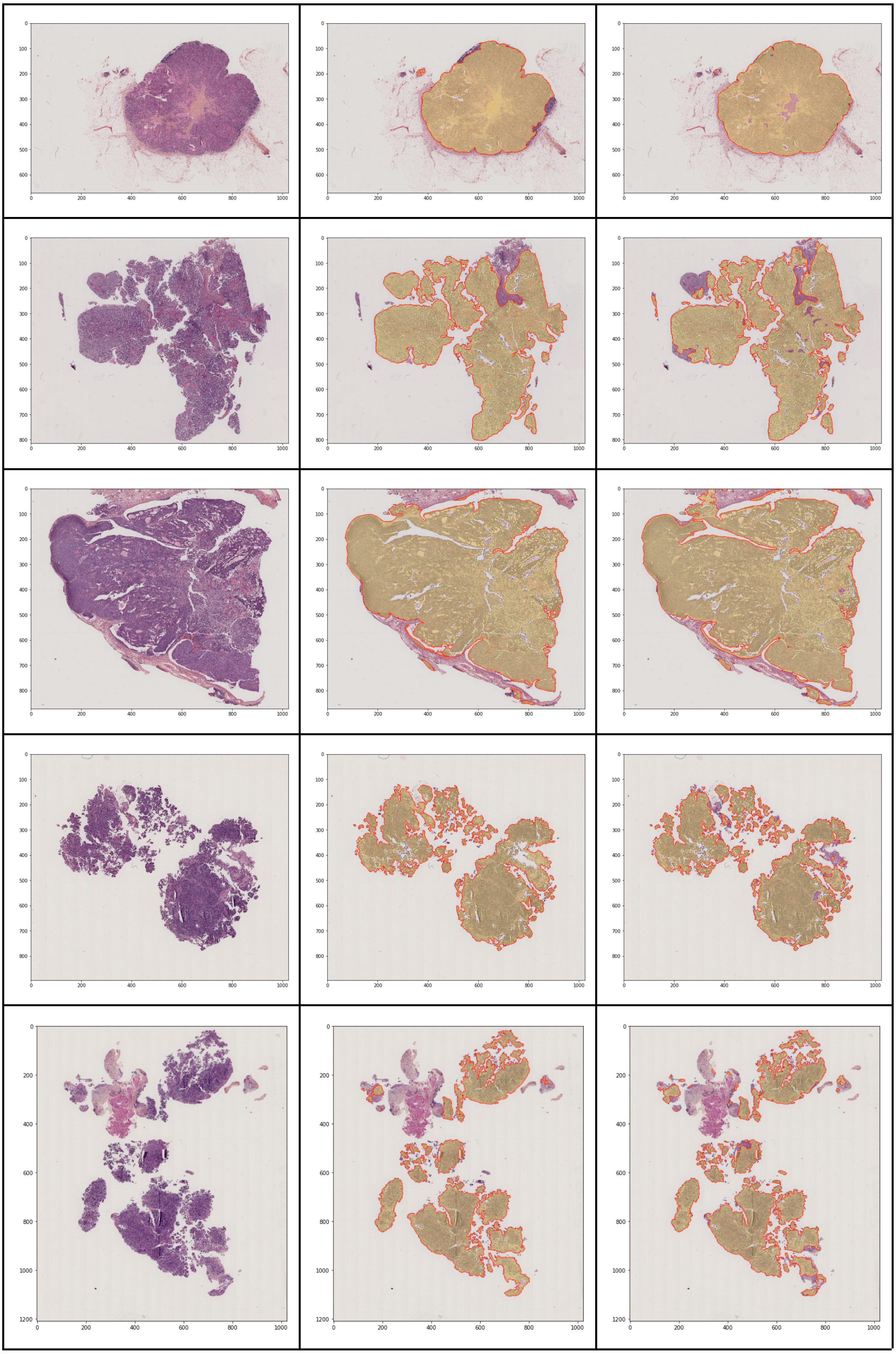

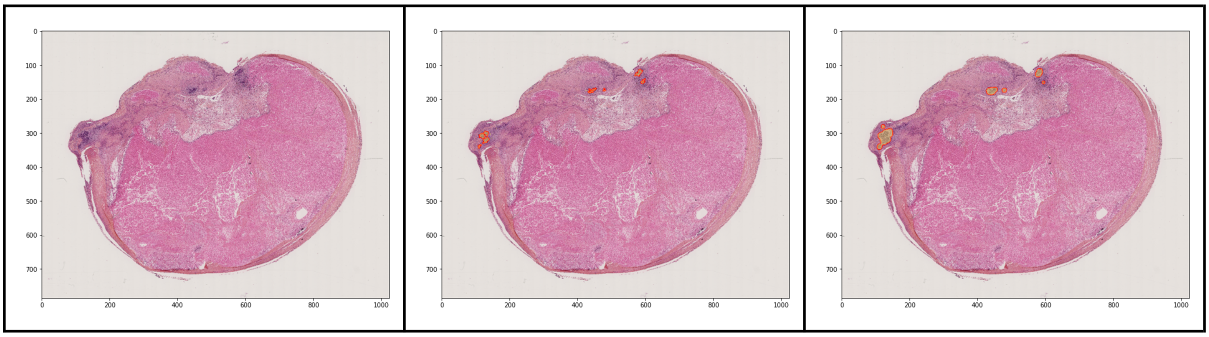
0
100
200
300
400
500
600
700
0
200
400
600
800
1000

**Supplementary figure 2**. WSI of Rennes delineated by the pathologist and the algorithm

| WSI | WSI delineated by the algorithm |
|---|---|
| 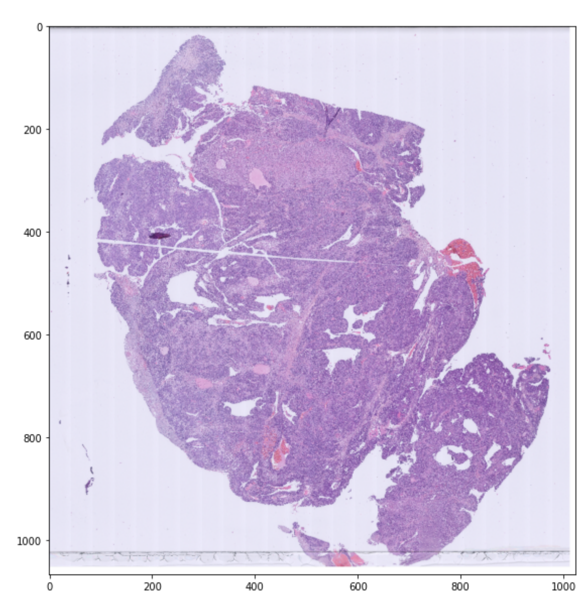 | 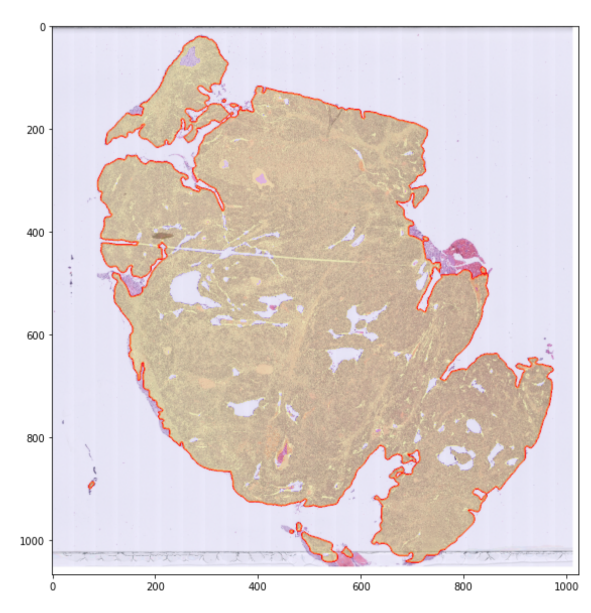 |
| 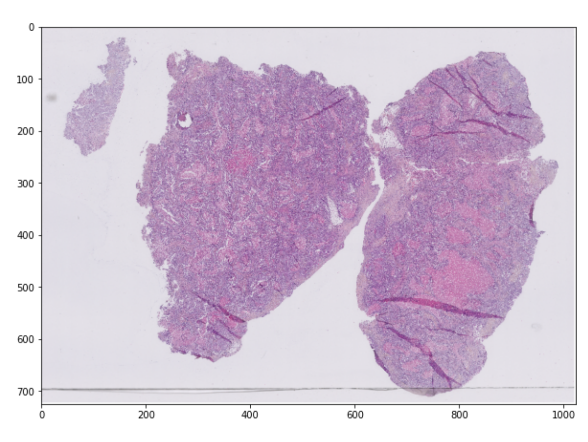 | 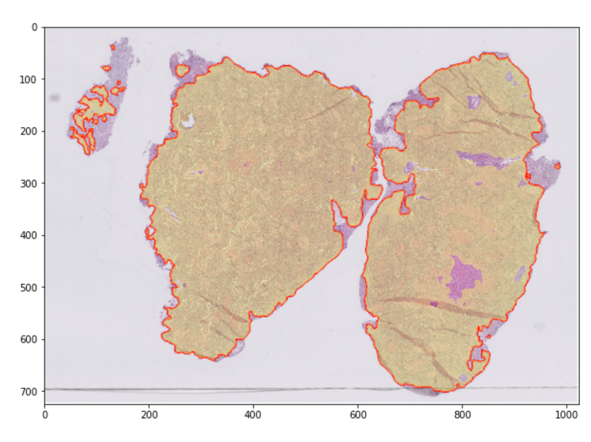 |
| 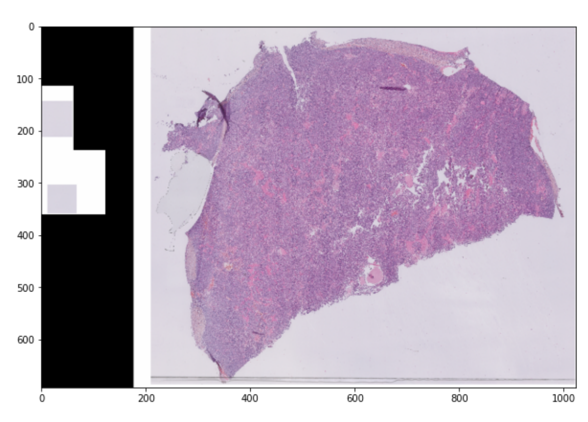 | 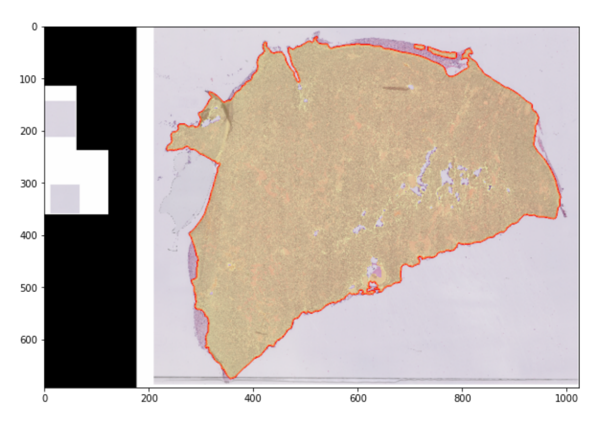 |

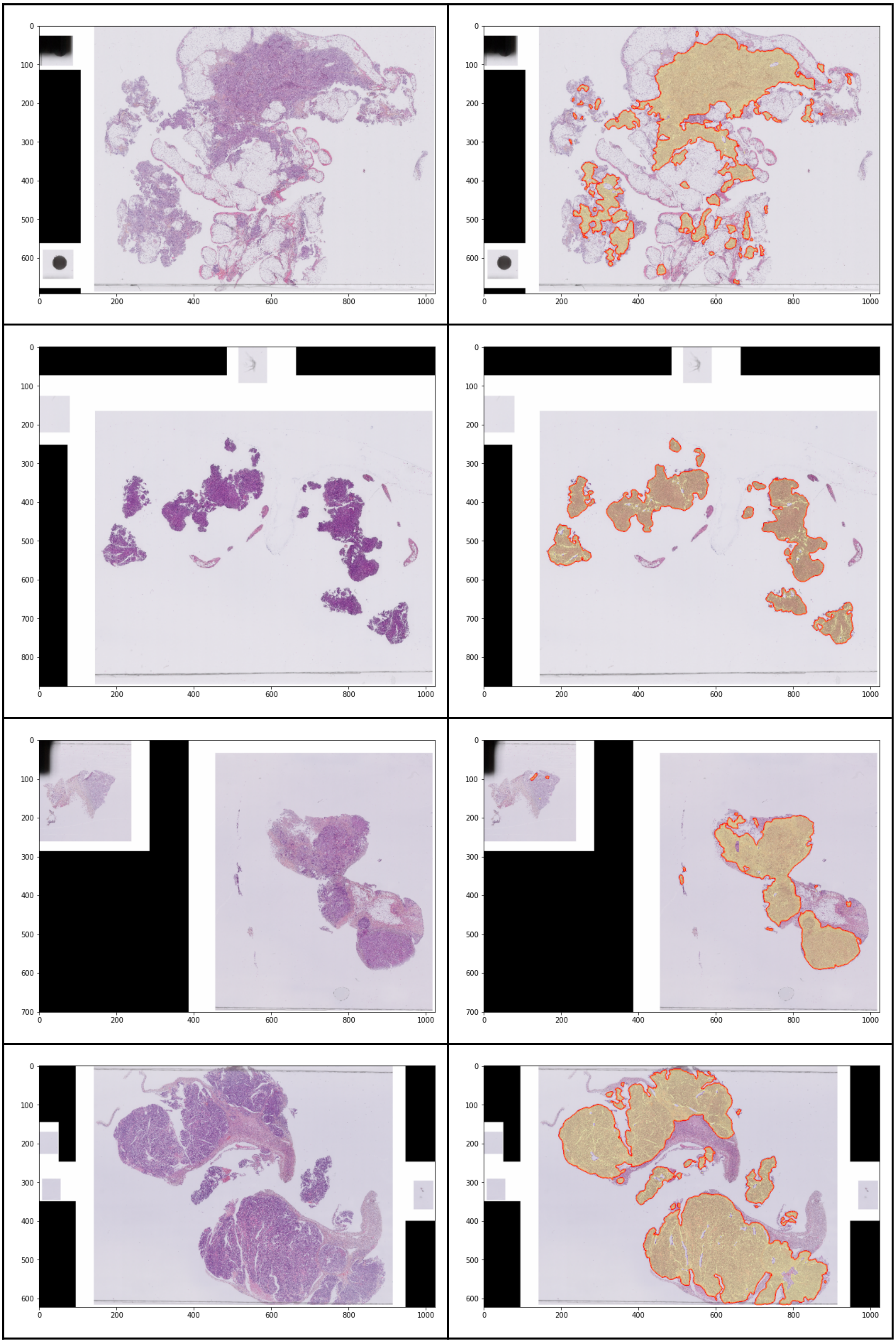
0 100 200 300 400 500 600
0 200 400 600 800 1000
0 100 200 300 400 500 600 700 800
0 200 400 600 800 1000
0 100 200 300 400 500 600 700
0 200 400 600 800 1000
0 100 200 300 400 500 600
0 200 400 600 800 1000

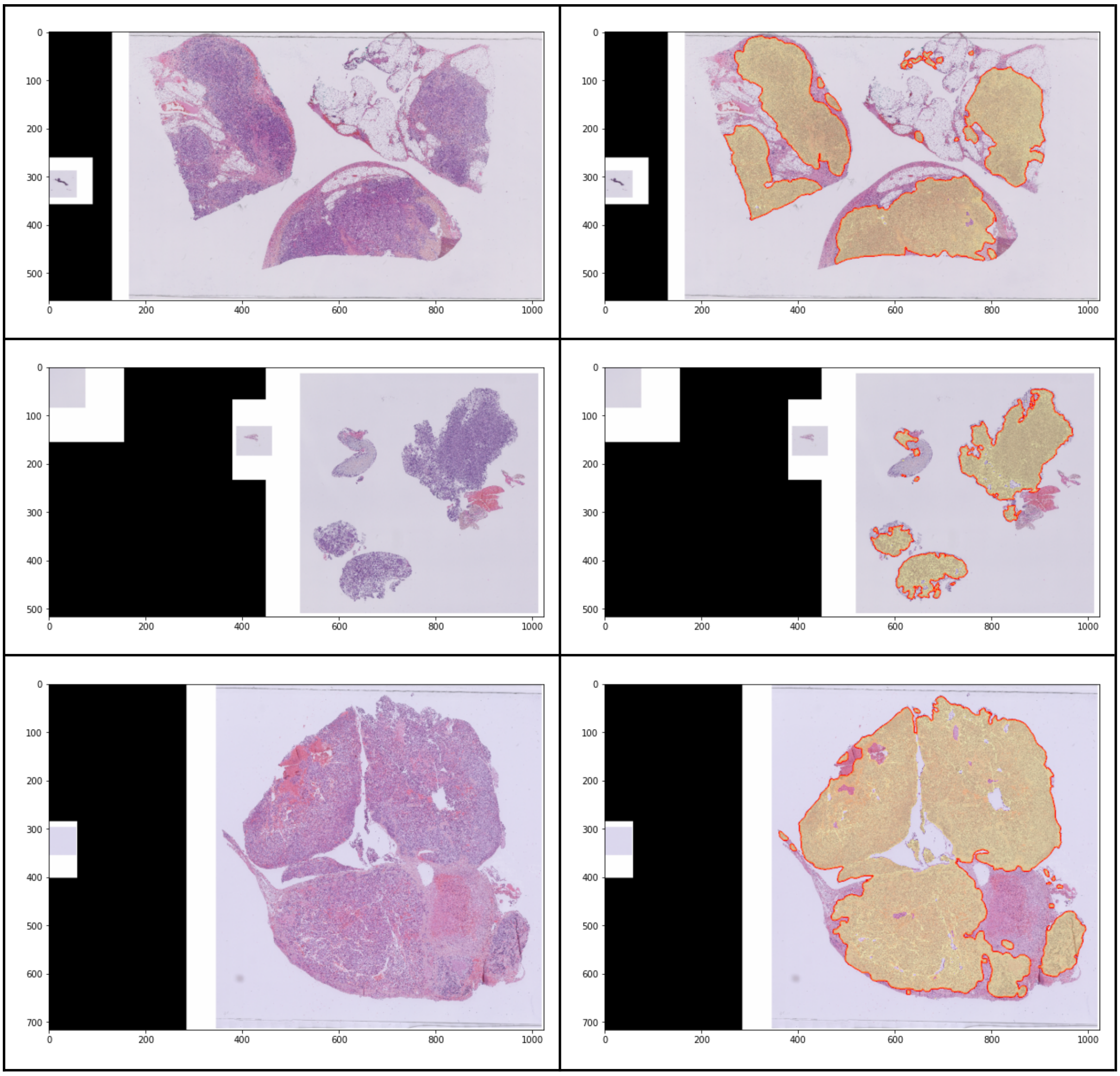
